# Low Temperatures Lead to Higher Toxicity of the Fungicide Folpet to Larval Stages of *Rana temporaria* and *Bufotes viridis*

**DOI:** 10.1101/2021.10.04.463096

**Authors:** Christoph Leeb, Laura Schuler, Carsten A. Brühl, Kathrin Theissinger

**Affiliations:** iES Landau, Institute for Environmental Sciences, University of Koblenz-Landau, Landau, Germany; LOEWE Centre for Translational Biodiversity Genomics, Senckenberg Biodiversity and Climate Research Centre, Frankfurt, Germany

**Keywords:** amphibia, climate change, green toad, common frog, pesticide, sensitivity, tadpole, temperature effect

## Abstract

Pesticides are one of the main drivers of the worldwide amphibian decline. Their actual toxicity depends on a number of factors, like the species in focus or the developmental stage of exposed individuals. As ectothermic species, the metabolism of amphibians is influenced by ambient temperature. Therefore, temperature also affects metabolic rates and thus processes that might enhance or reduce toxic effects. Studies about the interactive effect of temperature and toxicity on amphibians are rare and deliver contrasting results. To investigate the temperature-dependent pesticide sensitivity of larvae of two European species we conducted acute toxicity tests for the viticultural fungicide Folpan® 500 SC with the active ingredient folpet at different temperatures (6°C, 11°C, 16°C, 21°C, 26°C). Sensitivity of *Rana temporaria* and *Bufotes viridis* was highly affected by temperature: early larvae (Gosner stage 20) were about twice more sensitive to Folpan® 500 SC at 6°C compared to 21°C. Next to temperature, species and developmental stage of larvae had an effect on sensitivity. The most sensitive individuals (early stages of *R. temporaria* at 6°C) were 14.5 times more sensitive than the least sensitive ones (early stages of *B. viridis* at 26°C). Our results raise concerns about typical ecotoxicological studies with amphibians that are often conducted at temperatures between 15°C and 20°C. We suggest that future test designs should be performed at temperatures that reflect the temperature range amphibians are exposed to in their natural habitats. Variations in the sensitivity due to temperature should also be considered as an uncertainty factor in upcoming environmental risk assessments for amphibians.

## 1 Introduction

To improve crop yields about 360 million kg of pesticide formulations are used per year on agricultural fields in the European Union (data from 2017; Eurostat, 2020). Only a small part of these pesticides reaches their target organism [2], and due to spray drift and run-off they can get into water bodies within or near agricultural fields [3,4]. Such agricultural ponds can be important breeding habitats for amphibians [5–7], which are therefore exposed to pesticides during their aquatic life stages. Pesticides were shown to have adverse effects on amphibians in several studies (e.g. [8–12], and are consequently identified as one of the main drivers in the global amphibian decline [13,14]. The actual toxicity of pesticides for amphibians depends on a number of factors, including the active ingredients [8], formulation additives [9,10,15], the species in focus [12], a previous exposure to pesticides [10,11] and the developmental stage [10,12,16] of the tested individuals.

Also water temperature during pesticide exposure of larvae has an impact on the toxicity. Amphibians are ectothermic species and behavior and physiology are fundamentally influenced by environmental temperature [17]. Therefore, metabolic rates and thus processes that might enhance or reduce toxic effects, like the uptake of substances, the metabolic oxygen demand, and detoxification processes are temperature-dependent [18]. However, studies on the combined effects of temperature and pesticides on amphibians reveal contrasting results. Some observed that higher temperatures increased toxicity [19–21], while others showed a reduced toxic effect of pesticides on exposed amphibians [22–24]. For *Oligosoma polychroma*, a skink (reptile) and thus also an ectothermic vertebrate species, even a heat-seeking behavior was observed, that can be interpreted as response to increase the metabolism to better deal with stress after exposure to a glyphosate formulation [25].

Detailed knowledge of the relationship of pesticide sensitivity and temperature is central for two reasons. First, we are facing a global warming caused by climate change with more frequent temperature extremes [26]. Understanding the combined effect of this temperature increase and pesticides will help to better estimate the impact of climate change on amphibian populations, to identify potential threats on species and to set mitigation measures. Second, laboratory toxicity tests for pesticides with amphibian larvae are typically performed at temperatures between 15°C and 20°C (e.g. Mann et al., 2003; Johansson et al., 2006; Brühl et al., 2013; Wagner et al., 2017). These standard temperatures might not reflect the natural range of temperatures at which a species is exposed to pesticides in its habitat. For example, larvae of *Rana temporaria* can be found in European ponds with water temperatures only a few degrees above the freezing point [27]. In this study, the average water temperature during the aquatic development was 9.7°C and the maximum temperature 23°C [27]. However, in small water bodies the maximum water temperatures might be above 30°C, as even in high-altitudes breeding ponds with temperatures of up to 26.5°C can be found [28]. Therefore, standard laboratory toxicity tests might lead to the underestimation of possible sublethal or even lethal effects that occur at lower or higher temperatures. Thus, knowing the temperature at which amphibians are most sensitive will allow a more reliable assessment of the actual risk of pesticides.

In the present study, we conducted aquatic acute toxicity tests at temperatures between 6°C and 26°C to investigate the effect of the temperature on the sensitivity of amphibian larvae to the fungicide Folpan® 500 SC with the active ingredient folpet. With up to eight applications per growing season, folpet is, next to sulfur, the most common fungicide in German vineyards and is preventively used to protect plants primarily from mildew [29]. In general, fungicides are underrepresented in ecotoxicological studies compared to other pesticide classes [30]. To identify potential species and developmental stage specific differences in pesticide sensitivity, we tested early and late larval stages of the common frog (*Rana temporaria* Linnaeus, 1758) and the green toad (*Bufotes viridis* Laurenti, 1768), two temperate species that can be found in breeding ponds in German vineyards [6]. Both species are listed as “least concern” by the IUCN [31,32] and are widespread in Europe. *R. temporaria* is discussed as a model organism for European amphibian species in toxicological studies [33]. This species uses a variety of different water bodies for mating, which usually takes place in March, but can start as early as the end of January [34] when water temperatures are above 5°C for some days [34,35]. However, even at temperatures only a few degrees above freezing point spawning can be observed [36] and early larvae can be found [27]. Preferred temperatures of early *R. temporaria* larvae from Germany are between 14.8°C and 19.6°C, and between 16.5°C and 26.0°C of late larvae stages [37]. In contrast to *R. temporaria, B. viridis* is considered to be a thermophile species with preferred spawning temperatures between 16°C and 20°C. The optimum thermal tolerance limits for early larvae are between 12°C and 25°C [38].

The aim of the study was to get a better understanding about the temperature-dependent pesticide sensitivity of two European amphibian species. We hypothesized 1) that the sensitivity of larvae to Folpan® 500 SC is highly affected by water temperature, 2) that early larvae are more sensitive than late larval stages (see Adams and Brühl, 2020), and 3) that pesticide sensitivity differs between species.

## 2. Material and methods

### 2.1 Sampling and animal husbandry

Up to 300 eggs of eight and seven different clutches of *R. temporaria* and *B. viridis*, respectively, were collected in March and May 2018. The spawning pond of *R. temporaria* is located in the Palatinate Forest (Rhineland-Palatinate, Germany; 49.262433 N, 8.061896 E (WSG84), 242 m asl), distant from any pesticide use. The pond of the *B. viridis* population is located in a vineyard dominated area (Rhineland-Palatinate, Germany; 49.317490 N, 8.129091 E (WSG84), 194 m asl). Thus, the pond can be expected to be contaminated with various pesticides. Eggs were transferred to glass aquaria (30 × 20 × 20 cm) filled with tap water and kept in a climate chamber at 16°C with a 16:8 day-night-rhythm. For logistical reasons, not all acute toxicity tests for the same developmental stage were conducted at the same time. Therefore, parts of each clutch were kept at 10°C and daylight to slow down the development of the eggs. After hatching, larvae were kept in groups of 50 individuals in aerated glass aquaria filled with tap water at 21°C. As the larvae grew, we reduced their density to 20 larvae per aquaria. Cleaning of the aquaria and water renewal took place every second day. Larvae were fed daily *ad libitum* with commercial fish food, cooked salad, and cucumber.

### 2.2 Test substance

The fungicide Folpan® 500 SC (ADAMA Deutschland GmbH, Germany; purchased from a local distributor) with the active ingredient folpet (38-42% of weight; CAS number 133-07-03) was used for all tests. Folpet is an organochlorine phthalimide with a molecular weight of 296.6 g/mol and is used as a protective, broad-spectrum fungicide against leaf spot diseases in grapevines. Data on environmental contaminations are rare, but maximum measured concentrations of 50 ng/L in rivers [39] and 4.53 µg/L in ponds [40] have been reported. To assess the environmental realistic toxicity effect, the formulation was tested instead of the pure active ingredient. Other formulation ingredients are “alkylnaphthalensulfonic acid, polymer with formaldehyde, sodium salt” (3.5-5%), fumaric acid (1-1.5%), methenamine (0.5-1%) and 1,2-Benzisothiazoline-3-one (<0.1%). The acute aquatic toxicity of the formulation leads to a 96-h LC_50_ of 0.256 mg Folpan/L for the rainbow trout (*Oncorhynchus mykiss*) [41].

### 2.3 Experimental design

Acute toxicity of Folpan® 500 SC was determined in a full-factorial design with different temperature conditions and two developmental stages of both species. Early larval stages (Gosner stage 20; GS20; first hatchling stage with external gill circulation; see Gosner (1960) for classification) were tested at five different temperatures (6°C, 11°C, 16°C, 21°C, 26°C). Late larval stages (Gosner stage 36-41; GS40; larvae with at least hindlimbs) were tested at three different temperatures (6°C, 16°C, 26°C). For each combination of temperature, species and developmental stage (= 16 combinations in total), a 48 h static acute toxicity test was performed with six different pesticide concentrations, ranging between 0 (control) and 4.2 mg Folpan/L (see Supplementary Table 1). Fungicide concentrations were chosen based on range-finding tests and previous studies with folpet [16,43] to cover the concentration range at which ideally 0-100% mortality of the test organisms should be observed. Range finding tests were performed as 48 h tests with three Folpan concentrations and a control group with three replicates of one individual for each species/developmental stage and different temperatures. For each pesticide concentration of the final acute toxicity test 25 (GS20) or 15 (GS40) individuals were used, resulting in 150 and 90 individuals per test, respectively. Tests were conducted in 1.7 L glass jars containing 1L FETAX medium [44] and the respective amount of Folpan® 500 SC. Before adding the folpet formulation, the jars with the FETAX medium were cooled or heated to the test temperature in climate chambers (WK 19’/+15-35, Weiss Technik GmbH, Reiskirchen, Germany; MLR-351H SANYO Versatile Environmental Test Chamber, SANYO Electric Co. Ltd., Moriguchi, Japan). To reduce the influence of thermal shock on the physiology of the animals, preselected larvae from different clutches of about the same size and developmental stage (GS20 or GS40) showing normal behavior were placed in plastic boxes and acclimated at least for one hour to the test temperature. Afterwards five (GS20) or three (GS40) larvae were randomly placed in a test jar, resulting in five replicates/jars per pesticide concentration. For each jar, the mortality of larvae was determined after 48 h of exposure, whereby dead larvae were removed after 2 h and 24 h from the test jars. In accordance with the test guideline for acute toxicity testing in fish (OECD test guideline No. 203, [45]), larvae were not fed during the experimental period. Tests were performed in climate chambers set to the according test temperature with a 16:8 day-night-rhythm.

**Table 1:**
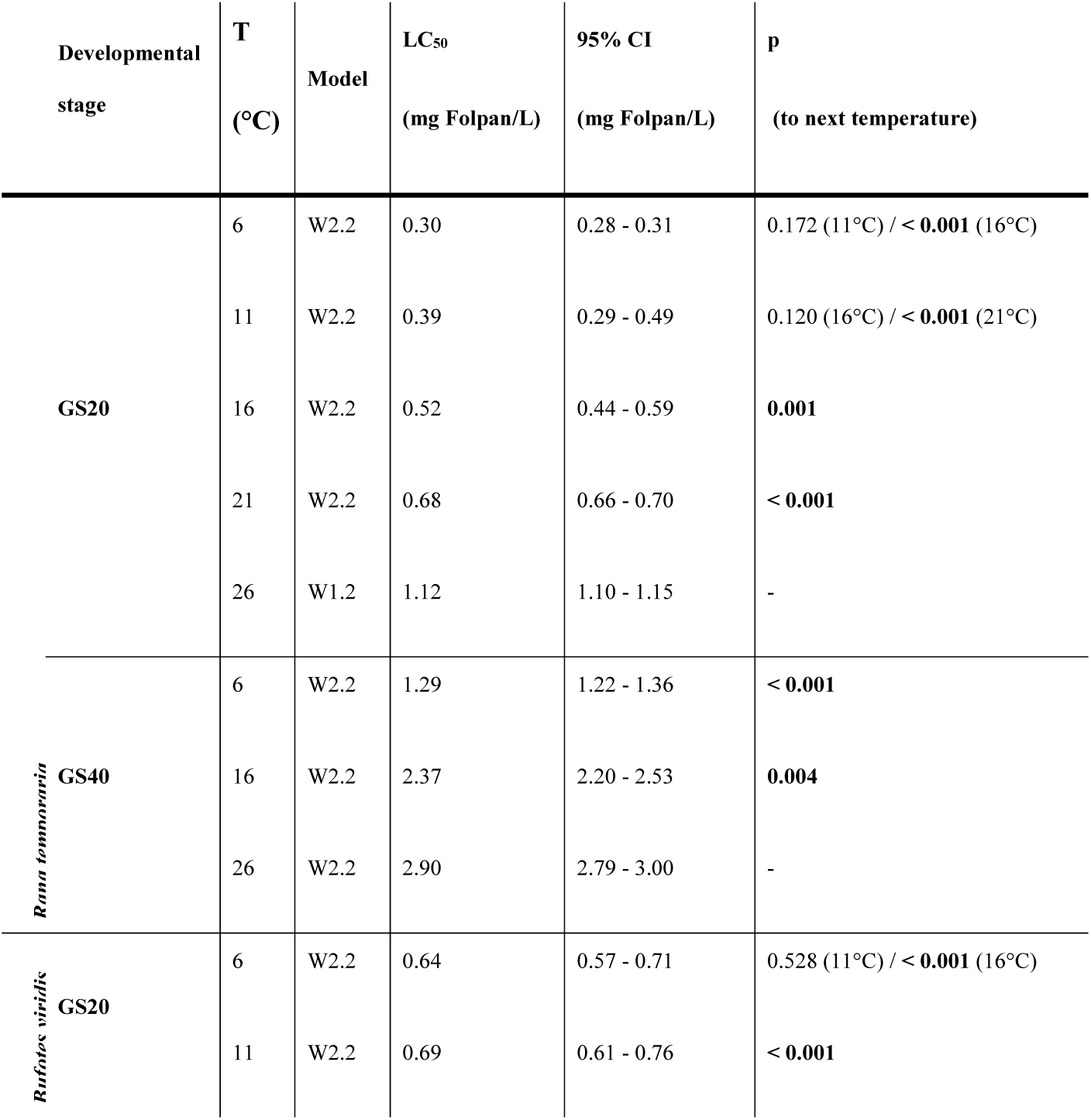
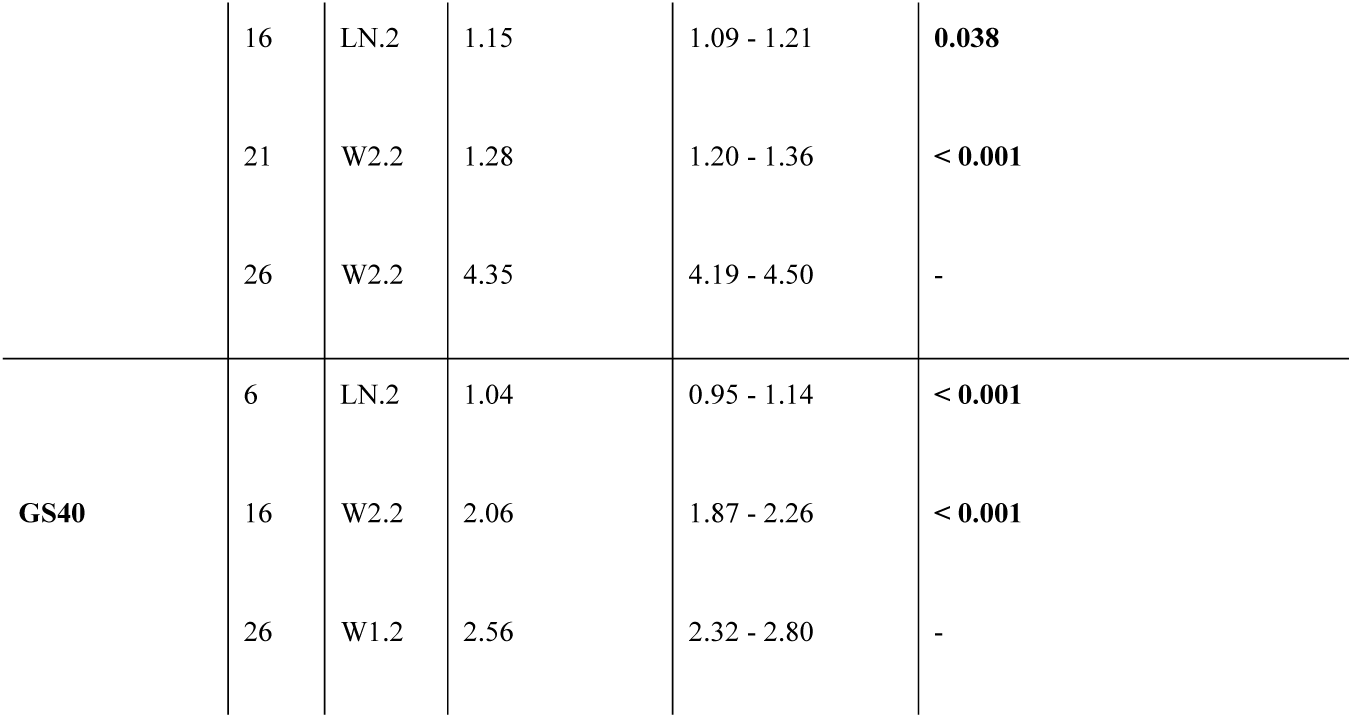
Calculated LC_50_ values for two developmental stages of *R. temporaria* and *B. viridis* at different temperatures with 95% confidence intervals and the used dose-response models. P-values show results from confidence interval overlap tests when testing against the next higher temperature. In case the difference was not significant, it was also tested against the two steps higher temperature. Significant differences after Bonferroni-correction are presented in bold.

### 2.4 Statistical analysis

For each test the median lethal concentration causing 50% mortality of test organisms (LC_50_ value) was determined using different concentration-response models (log-normal functions - LN.2, LN.3, LN.4; log-logistic functions - LL.2, LL.3u, LL.4, LL.5; and Weibull-functions - W1.2, W1.3, W1.4, W2.2, W2.3, W2.4) calculated with the R package “drc” [46]. To get the most accurate LC_50_ value, the model that best describes the observed mortality of larvae was selected based on the lowest Akaike’s Information Criterion for each test. LC_50_ values between different test temperatures for the same species and development stage were compared by a confidence interval overlap test [47] with the function “comped” implemented in “drc”. As we hypothesised a correlation between temperature and toxicity, we tested the LC_50_ of a species/developemental stage at a temperature only against the LC_50_ of the next higher temperature to reduce the probability of an alpha error accumulation. In case the difference was not significant, we also tested against the two steps higher temperature. Confidence interval overlap tests were also used to compare LC50 values between species and developement stages at the same test temperature. For all comparisons, p-values were calculated following the method described by Altman & Bland [48]. When testing the same species and developmental stage at different temperatures, or the same species or developemental stage at different temperatures, p-values were adjusted with a Bonferroni correction. All statistical analyses were carried out in R (version 3.4.3; R Core Team, 2019).

### 2.5 Animal welfare

The study was approved by the Landesuntersuchungsamt in Koblenz (Germany; approval number G18-20-009), and the collection of clutches and the husbandry of larvae were permitted by the “Struktur-und Genehmigungsdirektion Süd Referat 42 - Obere Naturschutzbehörde” (Neustadt an der Weinstraße, Germany; approval number: 42/553-254/455-18). After the experiments all test organisms were euthanized with a buffered 0.1% MS-222 solution.

## 3 Results

The calculated LC_50_ values of Folpan® 500 SC ranged between 0.30 and 2.90 mg Folpan/L for *R. temporaria* and 0.64 and 4.35 mg Folpan/L for *B. viridis* (Table 1). Toxicity decreased (i.e. increasing LC_50_ values) with increasing temperature for both tested species and developmental stages (see Fig. 1). In particular, the LC_50_ of GS20 at 21°C, the temperature at which toxicity tests are often conducted, was 2 (*R. temporaria*) and 2.3 (*B. viridis*) times higher than the lowest observed LC_50_ value. A temperature increase from 6°C to 16°C resulted in 1.7 to 2.0 and an increase from 16°C to 26°C in 1.2 to 3.8 times higher LC_50_ values. A temperature increase of 5°C (GS20) or 10°C (GS40) resulted always in a significantly higher LC_50_ value (all p ≤ 0.038, see Table 1), except for the comparison of 6°C and 11°C in GS20 in both species and 11°C and 16°C in GS20 *R. temporaria*. In general, the most sensitive individuals (*R. temporaria* GS20 at 6°C) were 14.5 times more sensitive than the least sensitive ones (*B. viridis* at GS20 26°C). Our analysis revealed that early larvae were more sensitive than late larvae, with the expection of *B. viridis* at 26°C (Table 2). Comparing LC_50_ values between species showed that *R. temporaria* is more sensitive in early and less sensitive in late developmental stages than *B. viridis* (Table 3), suggesting an interaction between developmental stage and species. However, the difference was not significant when comparing late developmental stages at 16°C and 26°C after a Bonferroni correction. Across all temperature treatments in both developmental stages and species the control and lowest concentration of 0.1 mg Folpan/L did not lead to any mortality in tested larvae.

**Table 2:** Comparison of LC_50_ values between developmental stages. Significant differences after Bonferroni-correction are presented in bold.

| Species | T (°C) | GS20 vs. GS40 |
| --- | --- | --- |
| *R. temporaria* | 6 | **< 0.001** |
|  | 16 | **< 0.001** |
|  | 26 | **< 0.001** |
| *B. viridis* | 6 | **< 0.001** |
|  | 16 | **< 0.001** |
|  | 26 | **< 0.001** |

**Table 3:** Comparison of LC_50_ values between species. Significant differences after Bonferroni-correction are presented in bold.

| Developmental stage | T (°C) | *R. temporaria* vs. *B. viridis* |
| --- | --- | --- |
| GS20 | 6 | **< 0.001** |
|  | 11 | **< 0.001** |
|  | 16 | **< 0.001** |
|  | 21 | **< 0.001** |
|  | 26 | **< 0.001** |
| GS40 | 6 | **0.002** |
|  | 16 | 0.088 |
|  | 26 | 0.063 |

**Figure 1:**
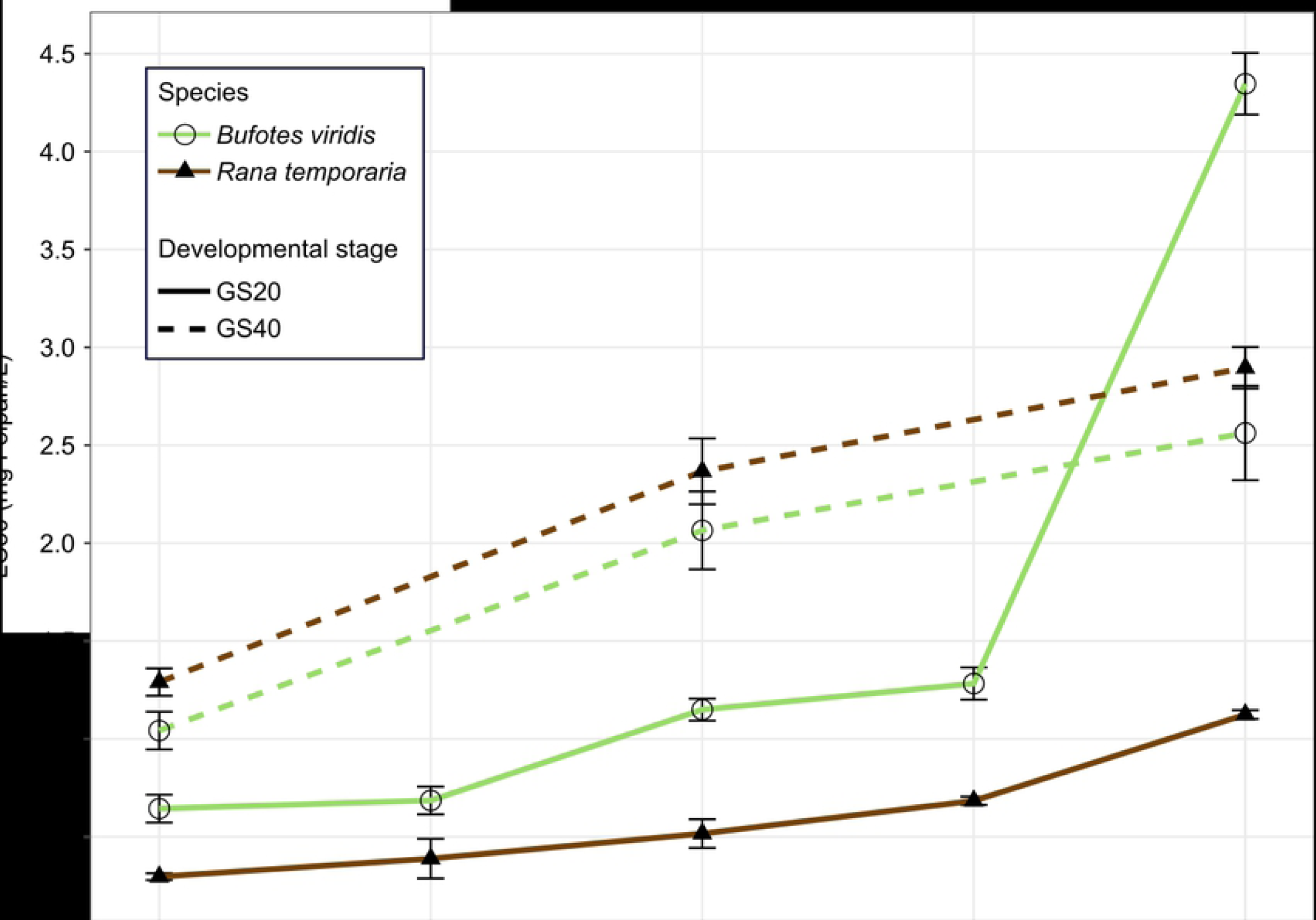
Calculated LC_50_ values (± 95% CI) of early (GS20) and late (GS40) developmental stages of *R. temporaria* and *B. viridis* at different temperatures. For detailed values and differences between temperatures see Table 1.

## 4 Discussion

In the present study, we demonstrated that the pesticide sensitivity of two European amphibian species is highly affected by temperature, with individuals of both tested developmental stages and species being more sensitive at lower temperatures. As we did not observe mortality at any temperature in controls, the tested temperatures are within a range that allows survival. Therefore, observed mortalities are caused by Folpan® 500 SC, where the lethal concentration depends on the temperature. Explanations for the relationship between temperature and sensitivity are diverse and depending on the pesticide and organism in focus, but exact mechanisms often remain unknown. In our study, higher temperatures might be nearer to the optimal temperature of the tested individuals, allowing effective metabolism and detoxification. Likewise, low temperatures might be below the optimal temperature range and result in additional stress, limiting the ability to cope with Folpan® 500 SC. Observed results might also be caused by the characteristics of folpet, the active ingredient of the tested formulation Folpan® 500 SC. In general, folpet degrades rapidly in aquatic environments and shows a half-life (DT50) of 0.7 h at 25°C and 0.178 h at 40°C (both pH 7; EFSA, 2009). Further, the degradation depends on the pH of the medium (DT50 pH 4, 25°C = 6.5 h; DT50 pH 4, 40°C = 1.06 h; DT50 ph 9, 25°C and 40°C = too rapid to measure; EFSA, 2009). Thus, the alkaline FETAX medium (ranging between pH 7.7 and 8.29 in our study) even accelerates the degradation. Although information about the degradation below 25°C is lacking, a temperature-dependent degradation that could have caused the observed effects can be expected. Because of the overall fast degradation, no analysis of the actual folpet concentration at the start and the end of a test was possible. It remains also unknown if the degradation of the formulation Folpan® 500 SC is similar to its active ingredient folpet, as additives could increase the stability of the formulation. Additives might also influence the toxicity of the formulation [9,10,15,51]. Regardless whether the lower sensitivity at higher temperatures is caused by a more effective metabolism and detoxification, and thus reduced bioaccumulation, or by an increased degeneration of folpet, Folpan® 500 SC is more toxic for the two tested amphibian species at lower temperatures.

Thus, increasing environmental temperatures might seem to have a positive effect on amphibians in terms of a reduced folpet toxicity. However, climate warming will also cause a shift in the breeding season to an earlier time of the year in temperate species [52]. Lötters et al. [53] showed that a shift of one month could decrease the glyphosate exposure risk during their migration to the breeding pond to about 50% for *R. temporaria*. Thus, also the exposure risk of larvae might be reduced. However, increased temperatures will also result in an earlier vegetation period of crops [54,55] and pesticides might be applied earlier. Consequently, the general exposure risk, but also the temperature at which amphibians will be exposed to pesticides in their aquatic habitats, will probably not change fundamentally. However, also more frequent temperature extremes can be expected [26], resulting in regional and temporary temperature drops so that also later larvae might be exposed to low temperatures. Climate change will also cause more frequent pesticide applications [56,57], resulting in higher overall pesticide loads in water bodies. Already today, many different pesticides can be found in ponds within agriculture [58,59]. Although higher temperatures might result in a lower sensitivity to folpet, contrary effects are possible for other pesticides and pesticide mixes. In vineyards, folpet is usually applied first in late May [60], when *R. temporaria* larvae occur in late development stages. At this time, *B. viridis* is still spawning and thus early larvae can be found. Only few data on actual environmental contamination with folpet are available and data show that maximum measured concentrations (50 ng/L [39]; 4.53 µg/L, [40]) are by a factor of at least 66 below the lowest LC_50_ values obtained in our study. We can therefore conclude that this pesticide will most likely not lethally affect the two tested amphibian species at the larval stage, but sublethal effects cannot be excluded. Thus, future studies should also focus on the effect of the temperature on sublethal endpoints like development or behavior.

Our results are in contrast to most studies that investigated the effect of temperature on pesticide toxicity for amphibian larvae in acute toxicity studies. In Materna et al. [20] leopard frog larvae (*Lithobates sp*.; former *R. pipiens* complex) showed higher mortalities in 96-h acute toxicity tests for the pyrethroid insecticide esfenvalerate at 22°C than at 18°C. Boone and Bridges [19] found the same relationship for *L. clamitans* (former *R. clamitans*) as the 96h-LC_50_ at 27°C was two times higher than at 17°C. Lau et al. [21] calculated 96h-LC_50_ values for the pesticide methomyl for three Asian amphibian species (*Duttaphrynus melanostictus, Polypedates megacephalus, Microhyla pulchra*) at temperatures between 15°C and 35°C, and observed lower 96h-LC_50_ values at higher temperatures. However, Chiari et al. [24] showed that increased temperature can also reduce the toxicity of a pesticide in 96-h acute toxicity tests by comparing published LC_50_ values for copper sulfate of various amphibian species. In contrast to most 96-h tests, reduced toxic effects of pesticides at higher temperatures can also be found in studies with tests running over several weeks or until metamorphosis. Baier et al. [23] found that the effects of the glyphosate formulation Roundup® PowerFlex on mortality, growth and tail deformation of the common toad (*Bufo bufo*) were more pronounced at 15°C than at 20°C. In a study on the glyphosate formulation Roundup® LB Plus, Baier et al. [22] also found increased effects on the development of common toad larvae at lower temperatures (15°C compared to 20°C) when exposure occurred already as egg. Rohr et al. [61] reported that an increased temperature reduced the time to the metamorphosis of larval *Ambystoma barbouri* exposed to the herbicide atrazine. Hence, also the total exposure to atrazine was reduced in this study, which ameliorated increased adverse effects of the pesticide [61].

With the exception of *B. viridis* at 26°C, early larval stages were 1.6 to 4.5 times more sensitive than late stages in both tested species. This is in line with the results from Adams and Brühl [16], where early larvae of *R. temporaria* (Gosner stage 20) were two times more sensitive than late larvae (Gosner stage 36) to the fungicide Folpan® 80 WDG with the same active ingredient folpet. Also Wagner et al. [10] found late larval stages of *R. temporaria* to be less sensitive in acute tests with two herbicides. Interestingly, in our study early larvae of *B. viridis* at 26°C were least sensitive. *Bufotes viridis* is a thermophilic species, and the highest tested temperature is at the upper limit of its optimal thermal range for development of early larvae (12°C - 25°C; Derakhshan and Nokhbatolfoghahai, 2015). Hence, 26°C might allow optimal detoxification without causing temperature stress for early stages. In late larval stages additional stress caused by processes linked to metamorphosis could countervail the advantages of high temperatures, resulting in late larvae of *B. viridis* being more sensitive than early larvae. Further, the optimal temperature of late *B. viridis* larvae could be even higher than 26°C. This might also explain why late larvae of *R. temporaria* (with assumed lower optimal temperature) were less sensitive than *B. viridis*, although *R. temporaria* is more sensitive in early stages. In general, species [12] and even population [10,11] specific differences in pesticide sensitivity are known. For example, Adams et al. [43] showed that out of eight central European amphibian species, the most sensitive species was five-times more sensitive than the least sensitive species towards the pesticide folpet. Therefore, differences in the sensitivity in our study species are not surprising. However, the original breeding pond where *B. viridis* eggs were obtained was situated within viticulture. Thus, it cannot be ruled out that differences in the sensitivity are the result of an adaption of the population to pesticides and not a species effect.

Folpet is, next to sulfur, the most common fungicide in German vineyards and thus understanding its toxicity on non-target organisms is of high relevance. However, the fast and temperature dependent degradation of folpet limits the conclusions drawn from our study. Thus, we recommend that future studies on the relationship of temperature and sensitivity of amphibians should focus on pesticides with a longer degradation time, not influenced as much by temperature. It might also be worth to consider pesticide mixtures, as often several formulations are applied at the same time [60] and a mixture of pesticides can be found in agricultural ponds [40]. It has recently been shown that the developmental temperature prior to ecotoxicological tests can have an influence on the organisms’ sensitivity to a test substance (Silva et al 2020) and should consequently also be considered in future amphibian tests.

To date, no standard test guideline for acute toxicity tests of European amphibian species exists and amphibians are also not explicitely considered in the environmntal risk assessment of pesticides. The results of our study raise concerns about typical ecotoxicological studies with amphibians that are often conducted at temperatures between 15°C and 20°C, because early larvae at 6°C were about two times more sensitive to Folpan® 500 SC as at 21°C. Therefore, adverse effects in temperate amphibian species might only be observed at lower or, depending on the tested pesticide, higher temperatures. Based on the results we obtained in our study we conclude that an additional temperature related factor needs to be incorporated in an uncertainty factor of an upcoming environmental risk assessments for amphibians in the EU that reflects variations in pesticide sensitivity due to temperature. Additionally, we agree with recommendations of previous studies [19,21–23] that future test protocols should consider temperature as an important factor. Tests should be performed at temperatures that are reflecting the temperature range amphibians are exposed to in their natural habitats, possibly also including natural daily temperature fluctuations.

## 6 Author contributions

CL, CB and KT conceived and designed the study. CL and LS performed the experiment. CL and LS analyzed the data and drafted the manuscript. KT acquired the funding of the project and supervised the work together with CB. All authors contributed to the writing process and approved the final manuscript.

## 8 Acknowledgments

We want to thank Elena Adams, Haiying Xia, Cedric Abele and Jana Huhle for their help in the laboratory.

## 9 Data availability statement

The original contributions presented in the study are included in the article/supplementary files, further inquiries can be directed to the corresponding author/s.

